# Overlapping local and systemic defense induced by an oomycete fatty acid MAMP and brown seaweed extract in tomato

**DOI:** 10.1101/2022.08.26.505481

**Authors:** Domonique C. Lewis, Danielle M. Stevens, Holly Little, Gitta L. Coaker, Richard M. Bostock

## Abstract

Eicosapolyenoic fatty acids are integral components of oomycete pathogens that can act as microbe-associated molecular patterns (MAMPs) to induce disease resistance in plants. Defense inducing eicosapolyenoic fatty acids include arachidonic (AA) and eicosapentaenoic acids, and are strong elicitors in solanaceous plants with bioactivity in other plant families. Similarly, extracts of the brown seaweed, *Ascophyllum nodosum*, used in sustainable agriculture as a biostimulant of plant growth, may also induce disease resistance. *A. nodosum*, similar to other macroalgae, is rich in eicosapolyenoic fatty acids, which comprise as much as 25% of total fatty acid composition. We investigated the response of roots and leaves from AA or a commercial *A. nodosum* extract (ANE) on root-treated tomatoes via RNA sequencing, phytohormone profiling, and disease assays. AA and ANE significantly altered transcriptional profiles relative to control plants, inducing numerous defense-related genes with both substantial overlap as well as differences in gene expression patterns. Root treatment with AA and, to a lesser extent, ANE also altered both salicylic acid and jasmonic acid levels while inducing local and systemic resistance to oomycete and bacterial pathogen challenge. Thus, our study highlights overlap in both local and systemic defense induced by AA and ANE, with potential for inducing broad-spectrum resistance against pathogens.

## Introduction

Members of Phaeophyta and Rhodophyta – brown and red macroalgae – contain multiple bioactive molecules and derived oligosaccharides that are known to induce defense responses in plants (Klarzynski et al., 2003; Sangha et al., 2010; Vera et al., 2011). *Ascophyllum nodosum*, a brown alga (seaweed), is a rich source of polyunsaturated fatty acids, including the eicosapolyenoic acids arachidonic acid (AA) and eicosapentaenoic acid (EPA), which comprise as much as 25% of total fatty acid composition (Lorenzo et al., 2017; van Ginneken et al., 2011). AA and EPA are essential fatty acids found in lipids and cell walls of oomycete pathogens, are absent from higher plants, and have specific structural requirements for elicitor activity (Araceli et al., 2007; Creamer and Bostock, 1986; Gellerman et al., 1975). Algal species like *A. nodosum* belong to the same major eukaryotic lineage as oomycetes, the Stramenopila. AA and EPA are potent oomycete-derived elicitors of plant defenses, and their elicitor activity is strongly enhanced by branched β-glucan oligosaccharins (Bostock et al., 1981; Robinson and Bostock, 2015). AA, EPA and other eicosapolyenoic acids (EPs) can be considered microbe-associated molecular patterns (MAMPs), although their initial perception and signaling is likely different than canonical MAMPs directly perceived by cell surface receptors (Ngou et al., 2022). EPs are released by *Phytophthora infestans* spores and hyphae during infection of potato leaves and are taken up and incorporated into host plant cell lipids or oxidized to hydroperoxy acids and uncharacterized products (Fournier et al., 1993; Göbel et al., 2002; Göbel et al., 2001; Hwang and Hwang, 2010; Preisig and Kuć, 1988; Ricker and Bostock, 1992; Véronési et al., 1996). Representing a novel class of MAMPs, AA and EPA engage hormone-mediated immune pathways in plants (Fidantsef and Bostock, 1998; Savchenko et al., 2010).

Extracts from *A. nodosum* are used in agriculture primarily to stimulate plant growth and development, but may also increase biotic and abiotic stress tolerances (Shukla et al., 2019). There are various commercial formulations of *A. nodosum* extracts, and each is a unique proprietary mixture. When compared, these products elicit varying transcriptional outcomes in plants (Goñi et al., 2016). The commercial *A. nodosum* extract, Acadian (hereafter ANE; APH-1011-00, Acadian Seaplants, Ltd., Nova Scotia, Canada), is a biostimulant that can also protect plants against fungal and bacterial pathogens (Ali et al., 2016; Jayaraj et al., 2008). In *Arabidopsis thaliana*, ANE induces systemic resistance to *Pseudomonas syringae* pv. *tomato* and *Sclerotinia sclerotiorum* (Subramanian et al., 2011). Studies on ANE-induced disease resistance in *A. thaliana* and tomato implicate jasmonic acid-dependent signaling, increased ROS production, induction of numerous immune response genes, and increased defense-related proteins and metabolites (Ali et al., 2016; Cook et al., 2018; Jayaraj et al., 2008; Subramanian et al., 2011). As the predominant polyunsaturated fatty acid in *A. nodosum*, AA may contribute to ANE’s biological activity. The ability of these EPs to induce resistance to diverse pathogens and trigger phytoalexin accumulation, lignin, reactive oxygen species, and programmed cell death has been shown in solanaceous and other plant families (Araceli et al., 2007; Bostock et al., 1981; Cook et al., 2018; Dye, 2020; Knight et al., 2001).

Here we investigate the overlap in plant response to AA and ANE. In this study, 3’ Batch Tag Sequencing (Lohman et al., 2016; Meyer et al., 2011) was used to compare and contrast AA- and ANE-induced transcriptomes locally in treated roots and systemically in leaves of root-treated tomato seedlings, revealing extensive overlap. Using disease assays with seedlings challenged with *Phytophthora capsici* and the bacterial speck pathogen, *Pseudomonas syringae* pv. *tomato* (*Pst*), we demonstrate the systemicity of AA- and ANE-induced resistance in tomato. The effect of AA and ANE root treatments on levels of selected phytohormones in roots and leaves also were determined to establish the relationship between transcriptional reprogramming and phytohormone changes that may prime or influence host defense.

## Results

### Transcriptomic analyses of AA- and ANE-induced plants

We hypothesized that AA- and ANE-induced resistance may be mediated by similar or shared transcriptional changes locally, in treated roots, and systemically, in leaves of root-treated tomato seedlings. To investigate transcriptomes of AA- and ANE-treated tomato (*Solanum lycopersicum cv*. ‘New Yorker’), roots of hydroponically grown seedlings were treated with 10 µM AA, 0.4 % ANE, 10 µM linoleic acid (LA), or H_2_O for 6, 24, and 48 hours prior to harvest and processing for RNA sequencing (**Fig. 1A**). Water and LA, an abundant fatty acid in plants, were included as negative controls. Of the sequencing reads across all samples, 65.3%-85.7% uniquely mapped to the tomato reference genome build SL 3.0 (**Sup. Fig 1**). Principle component analysis (PCA) of normalized read count data revealed distinct clustering of treatment groups across all tested timepoints in root tissue (**Fig. 1B-1D**). Both AA and ANE treatments exhibited unique clusters whereas control treatments, H_2_O and LA, clustered together at 6, 24 and 48 hours in roots. PCAs of read count data for leaf tissue showed similar but less distinct clustering across treatments and timepoints reflective of distal tissue (**Sup. Fig 2**). The most distinct clustering in leaves was observed at 24 hours where both negative controls, H_2_O and LA, overlapped. Partial overlap between AA and ANE treatment groups was also observed at 24 hours in leaf tissue. Similarly, heatmaps visualizing normalized read counts of the most differentially expressed (DE) genes by fold change at 24 hours showed distinct clustering by treatment group in both roots (**Fig. 2A**) and leaves (**Fig. 2B**). DE genes were set at an absolute fold change cut-off >4 and >2 for roots and leaves respectively with adjusted p-values < 0.05 across all timepoints and treatments. Heatmaps of transcriptomes depict clear grouping of profiles across both sampled tissues. Gene expression profiles of H_2_O and LA treatments were nearly indistinguishable but clearly distinct from the profiles resulting from AA and ANE treatments (**Fig. 1, Fig. 2**). AA and ANE induced robust transcriptional changes relative to control treatments (**Fig. 1, Fig. 2**), with both elicitors effecting significant overlap in gene expression profiles, as well as notable differences between treatments.

**Figure 1.**
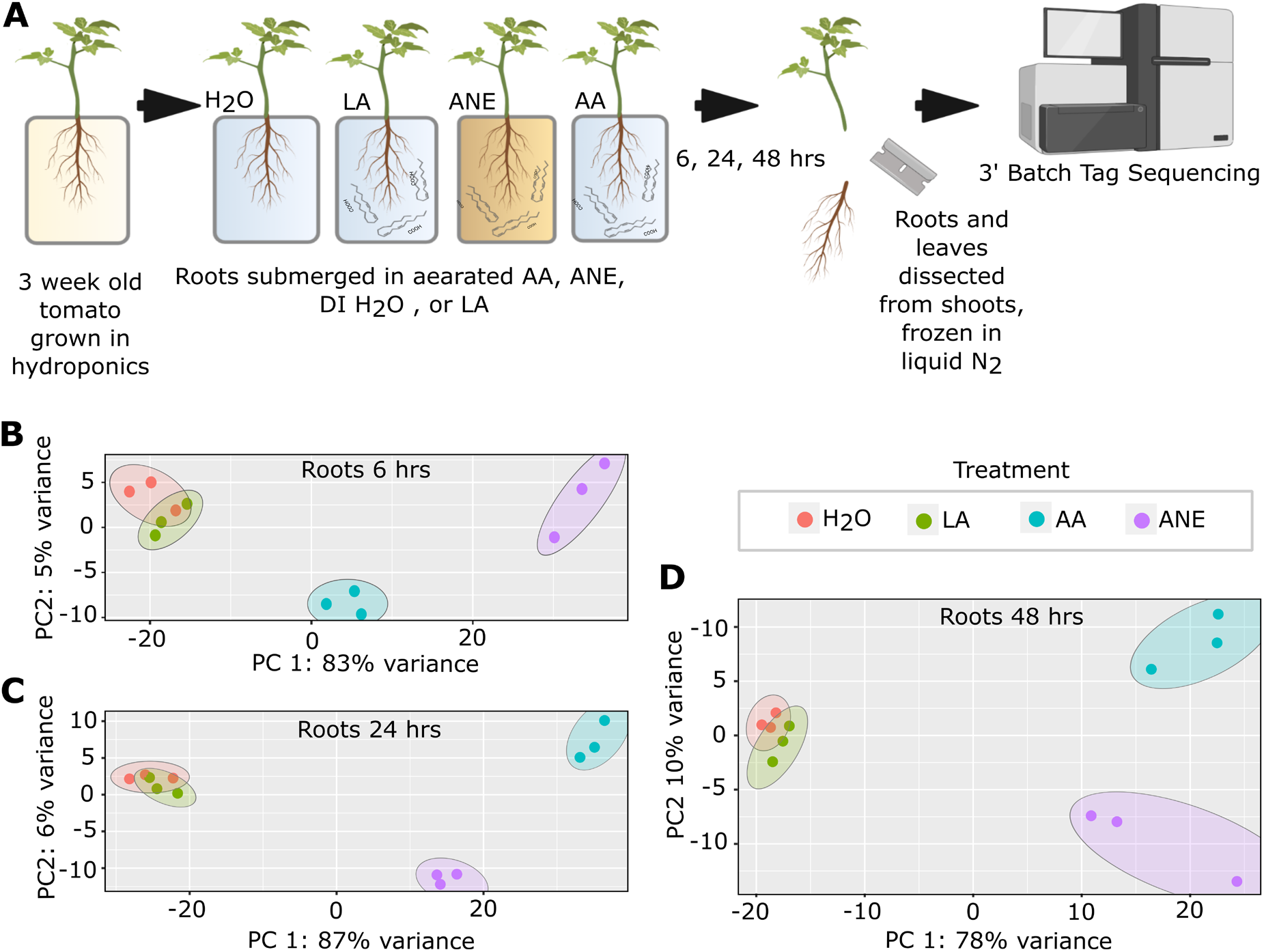
**(A)** Experimental procedure for RNA sequencing. Tomato roots treated with 10 µM Arachidonic Acid (AA), 0.4% Acadian (ANE), H_2_O, or 10 µM Linoleic acid (LA). Following 6, 24, and 48 hours root exposure to their respective treatments, plants were harvested, root and leaf tissue dissected from shoots, and the collected tissue flash frozen in liquid nitrogen. Harvested tissue was then subjected to total RNA extraction and DNase treatment. All samples were submitted for quality control, RNA-seq library construction and 3’-Batch-Tag-Sequencing. Principle component analysis scatterplots of RNA sequencing data in roots after **(B)** 6, **(C)** 24, and **(D)** 48 hours of treatment with 10 µM AA, 0.4% ANE, H_2_O, or 10 µM LA. PCA was conducted using the normalized read counts for all samples. PCA plots show variance of three biological replicates performed per timepoint and treatment.

**Figure 2.**
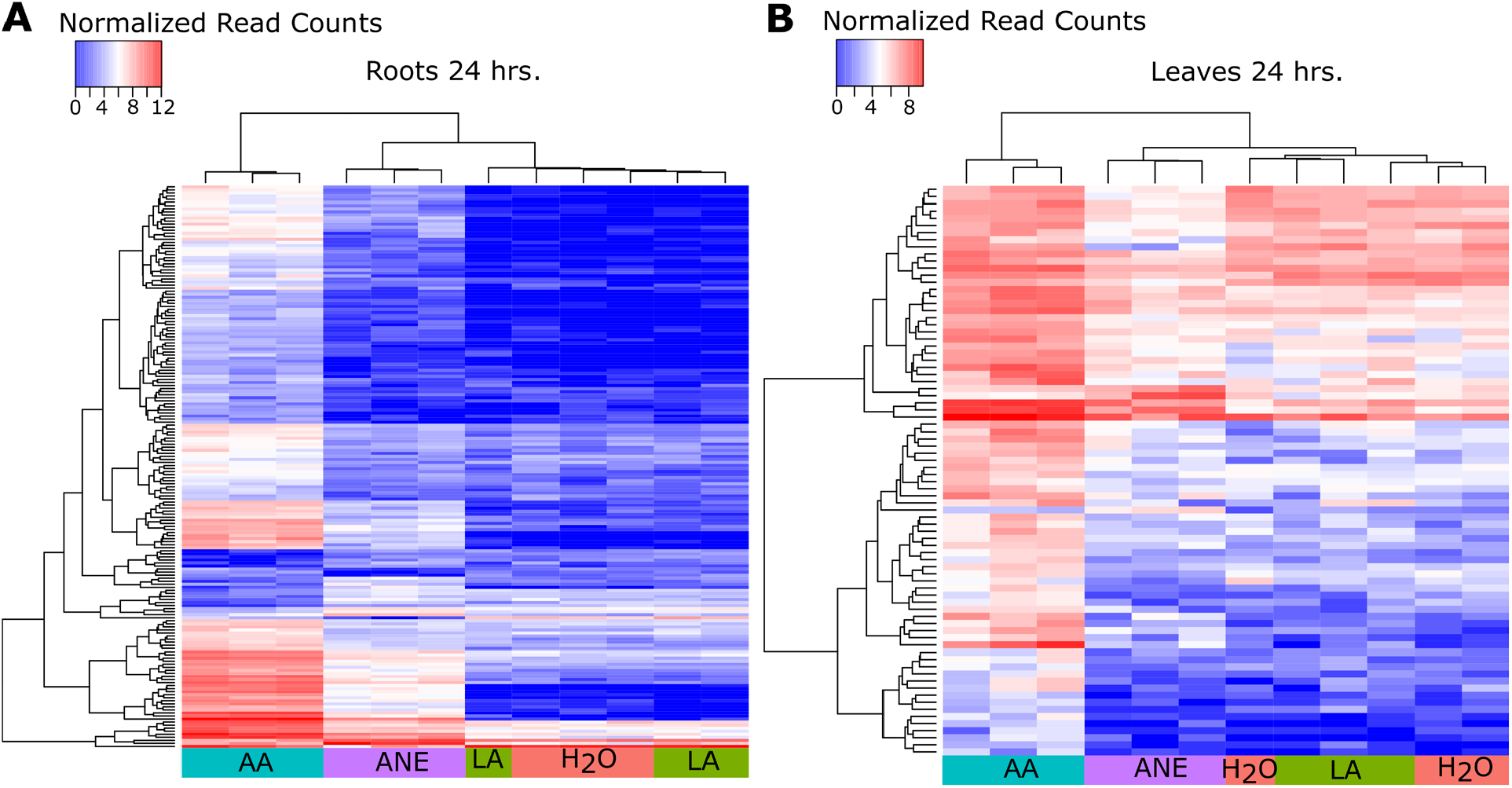
Normalized counts for the most differentially expressed genes by fold change for all treatment groups at 24 hours in **(A)** root and **(B)** leaf tissue. Plant roots were treated 10 µM Arachidonic Acid (AA), 0.4% Acadian (ANE), H_2_O, or 10 µM Linoleic acid (LA) for 24 hours. Blue indicates significant gene suppression and red indicated significant gene induction for each treatment. Heatmap data is log_2_ transformed and hierarchically clustered. Differentially expressed genes require an adjusted *P*-value < 0.05 and an absolute fold change in gene expression >4 for roots and >2 for leaves.

Root treatment with AA and ANE induced transcriptional changes relative to the H_2_O control both locally (roots) and systemically (leaves) with varying temporal dynamics. In AA-treated plants, transcriptional reprogramming occurred most strongly at 24 hours in roots and leaves (**Fig. 3A**). ANE-treated plants showed transcriptional changes most strongly at 6 hours in roots, and 24 hours in leaves (**Fig. 3A, Sup. Fig. 3**). Roots and leaves of either AA- or ANE-root-treated tomato seedlings have many shared DE genes, with AA-treated plants exhibiting the most numerous changes in gene expression. Within a tissue, roots and leaves shared up- and down-regulated DE genes for both AA and ANE treatments, with roots showing more DE genes than leaves (516 induced and 350 suppressed genes at 24 hours; **Fig. 3C**). By comparison, leaves had 51 induced and 77 suppressed genes at 24 hours (**Sup. Fig. 3**).

**Figure 3.**
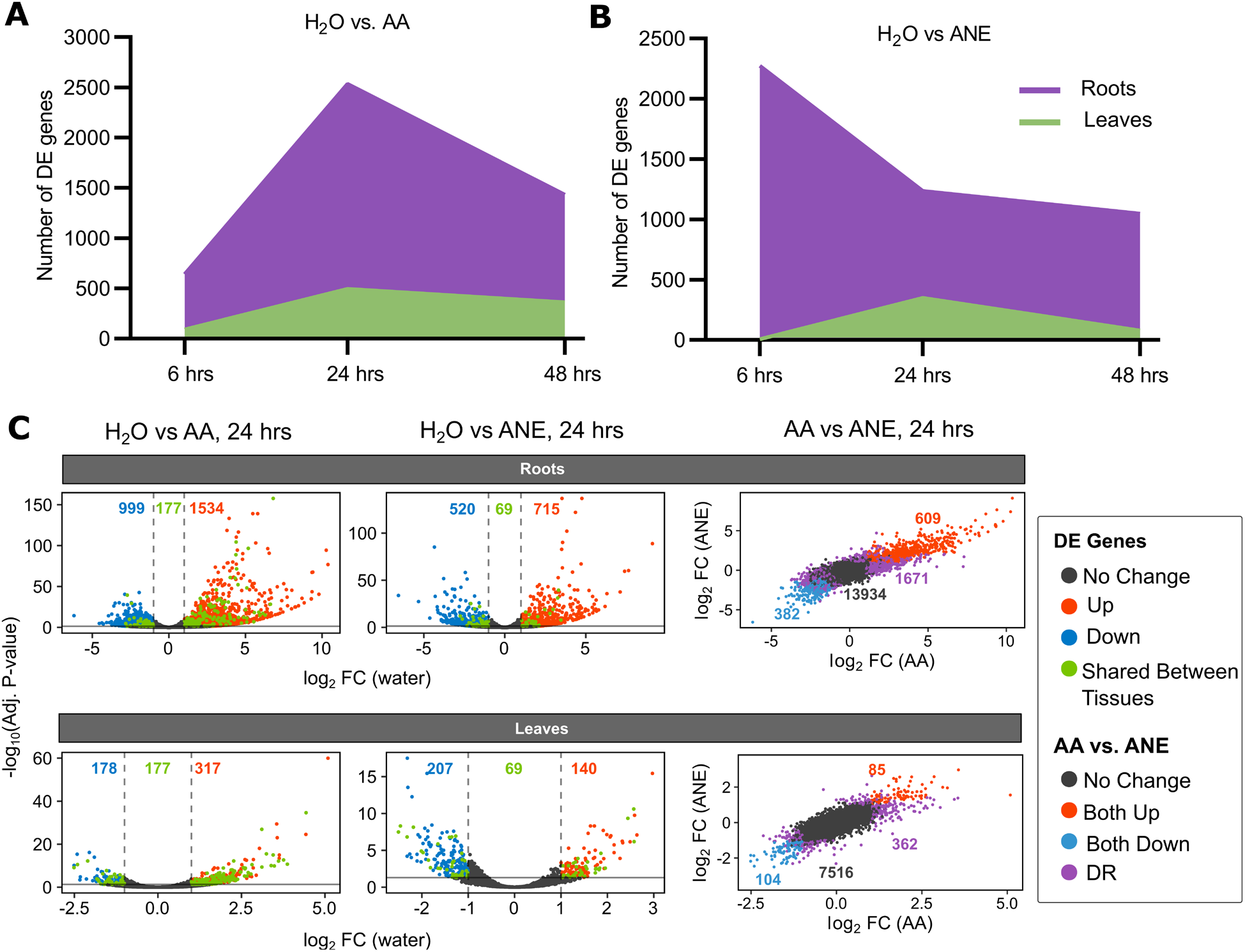
Total number of significant differentially expressed (DE) genes compared to H_2_O by tissue across time points for **(A)** Arachidonic Acid (AA) and **(B)** Acadian (ANE) root-treated tomato. Plant roots were treated with H_2_O, 10 µM LA, 10 µM AA, or 0.4% ANE for 6, 24 or 48 hours. **(C) Left:** Scatter plot of DE genes with the number of significantly up (red) and down (blue) regulated genes plotted at 24 hours from roots and leaves treated with AA or ANE compared to H_2_O. Those DE genes shared between tissues within a treatment are colored green. Solid and dashed lines represent cutoffs for significant DE genes (adjusted *P*-value < 0.05 and an absolute fold change in gene expression >2). **Right:** Scatter plot of gene expression in response to AA vs. ANE within the same tissue colored by differential response: both no change (grey), up regulated (red), and down regulated (blue) using the same significant cutoffs. Genes with different response between treatments are labeled DR (purple).

Although AA and ANE root treatments altered expression of many of the same genes, these treatments also induced distinct transcriptome features in roots and leaves. Of genes that were unique to each treatment, roots displayed a higher number of these features with 1585 genes identified compared to 263 genes in leaves (**Fig. 3C**). At the earliest tested timepoint, the transcriptional profile of ANE-treated roots revealed more than 76% unique DE genes. At 24- and 48-hour time points, AA- and ANE-treated roots showed the most overlap in transcriptional changes with some 992 and 728 shared DE genes, respectively. Analysis of distal untreated leaf tissue also revealed a similar trend with overlap in shared DE genes occurring most robustly at 24 and 48 hours (**Sup. Fig. 3D**). Distinct transcriptional features in the leaves of AA and ANE root-treated plants can be seen across all tested timepoints with more than 61% and 45% of identified DE genes being specific to their respective treatment group at 24 hours (**Sup. Fig. 3**).

### AA and ANE treatment induce upregulation of transcripts involved in oxylipins, immunity, and secondary metabolism

In order to identify specific gene categories and biological processes altered by AA and ANE treatments, we performed gene ontology (GO) functional analyses. GO analyses revealed AA and ANE root treatments enrich similar categories of tomato genes in both molecular function and biological processes categories **(Fig. 4A-4B)**. AA and ANE enriched root transcripts associated with oxidation-reduction processes, including hydrogen peroxide catabolism, oxidative stress responses, and heme binding. Both treatments induced cell wall macromolecule catabolism genes, as well as a variety of genes classically associated with defense responses.

**Figure 4.**
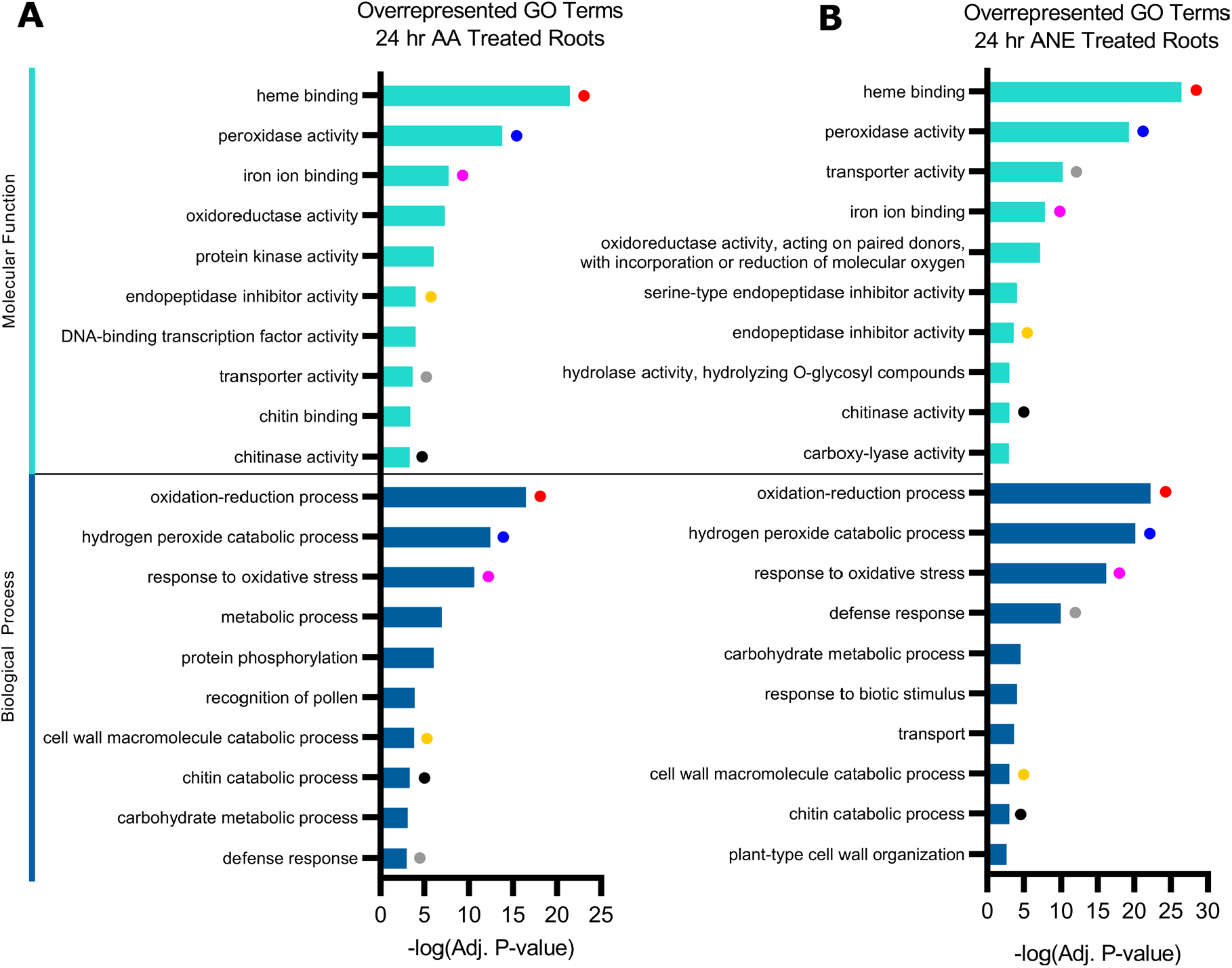
Gene ontology (GO) functional analysis of DE genes in roots 24 hours post-treatment. GO enrichment was conducted using Goseq. The top 10 most significantly (*P*-value < 0.05) enriched GO terms in molecular function and biological process GO categories are shown. Colored dots indicate shared molecular function and biological processes between **(A)** Arachidonic Acid (AA) and **(B)** Acadian (ANE) treatments. All adjusted *P*-values are negative 10-base log transformed.

Identification of specific defense- and stress-related genes significantly induced in AA- and ANE-treated roots at 24 hours revealed insightful overlap (**Fig. 7**). Transcripts of several key genes in biosynthesis of plant oxylipins including *α-DOX1, 9-DES, 9-LOX*, and *AOS3* were significantly up-regulated in response to AA and ANE root treatment. This corresponds with early work that first implicated oxylipin metabolism in AA action and demonstrated the capacity of plant endogenous *9-LOX* to use AA as a substrate (Andreou et al., 2009; Fournier et al., 1993; Göbel et al., 2002; Göbel et al., 2001; Hwang and Hwang, 2010; Véronési et al., 1996). The phytohormone jasmonic acid (JA) and related metabolites are also oxylipins (Dave and Graham, 2012). We observed induction of multiple JA-responsive genes including *JA2, JA2L, JAZ11*, and *JAZ7*, which previous studies reported induction in response to *Pst* infection or exogenous application of various phytohormones (Chini et al., 2017; Du et al., 2014; Ishiga et al., 2013).

AA and ANE similarly induced genes encoding known immune signaling components in roots at 24 hours. Significant increase in expression was seen in genes encoding MAPKKK’s and several WRKY transcription factors, including *WRKY39*, which confers enhanced resistance to biotic and abiotic stressors upon overexpression in transgenic tomato (Sun et al., 2015). Accumulation of salicylic acid (SA) and induction of SA-responsive genes are hallmarks of plant immune responses, including MAMP perception (Chen et al., 2017; Tsuda et al., 2009). In roots at 24 hours we observed upregulation of *NPR1* (**Fig. 7**), encoding a SA receptor that positively regulates expression of SA-dependent genes and is considered a master regulator of SA signaling (Chen et al., 2017). Likewise, the SA marker and pathogenesis-related (PR) gene *PR-1* showed induction in roots at 24 hours. Shared concurrent induction of these immunity related genes indicate plants exposed to AA and ANE are generally primed for defense against wide array of potential pathogen challenge.

Genes involved in secondary metabolism also showed strong induction in roots at 24 hours. Shikimate pathway members phenylalanine ammonia-lyase (*PAL*) and chorismate synthase, *CS1*, had increased expression compared to water in AA- and ANE-treated roots. Up-regulation of genes involved in metabolism of other phenolic compounds included tyramine n-hydroxycinnamoyl transferase (*THT1-3*) and a polyphenol oxidase. The sesquiterpenoid biosynthesis gene viridiflorene synthase, *TSP31*, showed significant induction with an increase in log_2_ fold change of 9.29 and 5.66 for AA and ANE, respectively, compared to water. Genes for key early steps in terpenoid biosynthesis, 3-hydroxy-3-methylglutaryl coenzyme A synthase (*HMGCS*) and 3-hydroxy-3-methylglutaryl coenzyme A reductase 2 (*HMGCR2*), a pathogen- and elicitor-responsive isoform, showed robust increases in expression in both treatment groups (Choi et al., 1992; Stermer and Bostock, 1987).

To assess global trends in transcriptional remodeling, GO functional analyses of AA- and ANE-treated tomato revealed significant congruency in under-represented gene categories. Nearly perfect overlap was seen in all unenriched GO terms in molecular function, biological process, and cellular compartment in roots at 24 hours (**Sup. Table 1**). Examination of specific shared genes most strongly down-regulated in response to AA and ANE treatment revealed suppression of genes associated with metal transport (**Sup. Table 2**). This included genes annotated to operate as metal, iron-regulated and copper transporters and metal tolerance in roots at 24 hours. The uptake and translocation of nutrient metals is essential for plant growth and development (Jogawat et al., 2021). The down-regulation of these transporters may indicate a shift toward defense rather than growth in the plant.

### Transcriptional changes specific to AA and to ANE treatments

Considering the difference in composition of AA and ANE, we examined the strongest uniquely up- and down-regulated genes in roots at 24 hours (**Sup. Table 3**). Unique transcriptional responses for AA-treated plants revealed differential expression of ethylene and terpene biosynthesis genes and modulation of genes involved in auxin signaling, cell wall anabolism, and signaling peptide formation. This included significant induction of 1-aminocyclopropane-1-carboxylate synthase 2 (*ACS2*), and suppression of an *ACO* isoform encoding 1-aminocyclopropane-1-carboxylate oxidase-like protein. Unique up regulation was also seen in a purported sesquiterpene synthase gene that showed a log_2_ fold change of 5.02 compared to H_2_O. Unique differential gene expression after AA treatment was also observed in small auxin up-regulated *RNA 36* (*SAUR36*) and a gene encoding a purported auxin efflux carrier. AA-treated plants showed induction of *PKS3L* which encodes a precursor of the immune signaling peptide, phytosulfokine (PKS), a recently classified damage associated molecular pattern (Zhang et al., 2018).

Unique transcriptional responses for ANE-treated roots at 24 hours include those involved in auxin signaling, cytokinin biosynthesis, specialized plant metabolism, cell proliferation, and induced resistance. This included robust induction of *SAR8*.*2*, encoding a systemic acquired resistance protein, and *CKX2*, encoding cytokinin oxidase 2. Significant induction was also observed for *IAA2*, an auxin regulated transcription factor. ANE-treated plants also showed up-regulation of *TCMP-1*, a tomato metallocarboxypeptidase inhibitor. Modulation was also seen in two 2-oxoglutarate and Fe(II)-dependent oxygenase superfamily members (*2ODD*s). These data collectively show AA and ANE locally and systemically alter the transcriptional profile of tomato through modulation of many defense-related genes.

### Phytohormone Quantification

AA and ANE modulate expression of genes associated with JA, SA, and ethylene phytohormone signaling and biosynthesis. Therefore, levels of selected phytohormones and phytohormone precursors were quantified via LC-MS/MS (**Fig. 5**). The JA precursor, oxophytodienoic acid (OPDA), accumulated in the leaves of AA- and ANE-root-treated plants at 48 hours, while accumulation of JA and its isoleucine conjugate (JA-Ile) occurred in the roots of AA-treated roots at 24 hours. This coincides with induction of JA signaling components *JA2, JA2L, JAZ7*, and *JAZ11* in the roots of AA-treated plants at 24 hours (**Table 1**). Salicylic acid (SA) accumulated in roots of AA-treated plants at both sample time points, consistent with our observation of transcriptional upregulation of *NPR1* and *PR1* (**Table 1**). Elevated levels of SA also were seen in leaves of both AA- and ANE-root-treated seedlings at 48 hours. Abscisic acid (ABA) increased in leaf tissue 24 and 48 hours after root treatment with AA or ANE. Indole acetic acid (IAA) and its aspartate conjugate (IAA-Asp) and zeatin riboside isomers were reduced at 24 hours in the roots of both AA- and ANE-treated plants. These changes in accumulation of IAA and its conjugate are consistent with the unique differential gene expression in small auxin up-regulated *RNA 36* (*SAUR36*) and a gene encoding a purported auxin efflux carrier in AA-treated roots at 24 hours. Likewise altered levels of IAA/IAA-Asp also coincide with the unique induction of *IAA2*, an auxin-regulated transcription factor, in the roots of ANE-treated plants at 24 hours. The leaves of AA- and ANE-root-treated plants also had reduced levels of zeatin ribosides at 24 and 48 hours. Taken together, these data demonstrate that AA and ANE both alter the accumulation of multiple phytohormones, including those that modulate defense networks in tomato.

**Figure 5.**
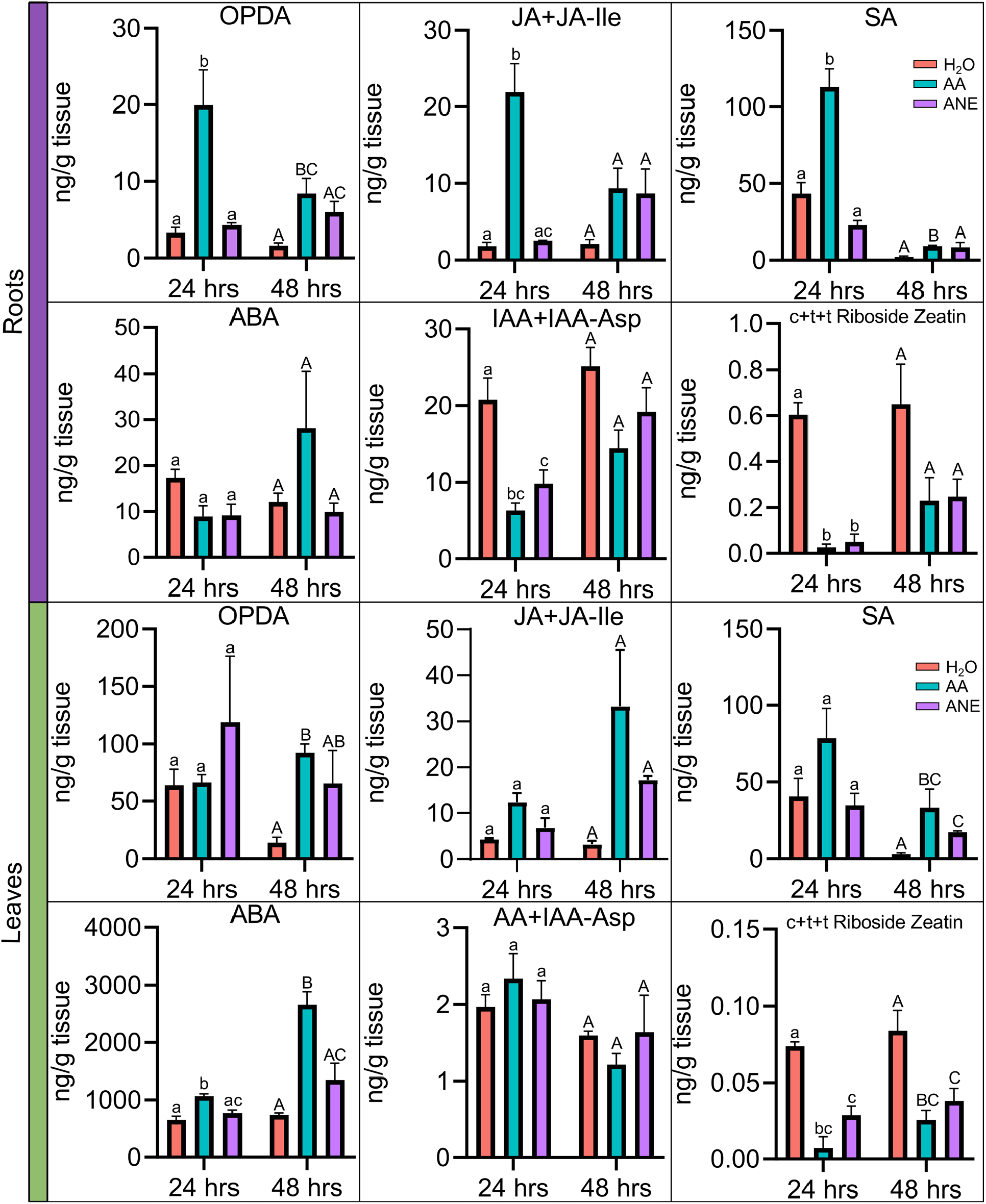
Quantification of phytohormones oxophytodienoic acid (OPDA), Jasmonic acid and JA-isoleucine (JA+JA-Ile), salicylic acid (SA), abscisic acid (ABA), indole acetic acid and IAA-aspartate (IAA+IAA-Asp), cis/trans zeatin and trans Riboside zeatin in **(A)** roots and **(B)** leaves of tomato seedlings root-treated with H_2_O, 10 µM AA, or 0.4% ANE, for 24 and 48 hours. Error bars represent standard error of 3 biological replicates. For bars with different letters, the difference of means is statistically significant by ANOVA and Tukey’s HSD *P*<0.05. Lower-case letters denote statistical significance for 24 hours and upper-case letters denote statistical significance for 48 hours. All statistical comparisons are within a single timepoint and tissue type.

### Local and systemic induced resistance

Given the overlapping transcriptional profiles and clear changes in phytohormone accumulation induced by AA and ANE root treatment, we utilized disease assays to establish the systemic nature of AA-induced resistance and to investigate the local and systemic nature of ANE-induced resistance. AA-, ANE-, or H_2_O-pretreated roots were inoculated with zoospores of the oomycete *P. capsici* and then seedlings were evaluated for collapse due to root and crown rot. Post-inoculation, 85% of plants treated with H_2_O collapsed at the crown while less than 20% of ANE-treated plants collapsed. Treatment of roots with 0.4% ANE protected seedlings against *Phytophthora* root and crown rot compared to H_2_O-treated/inoculated control seedlings (**Fig. 6A and C**). ANE’s level of protection is similar to that observed with AA-induced resistance in tomato and pepper to *P. capsici* infection using this same assay format (Dye and Bostock, 2021).

**Figure 6.**
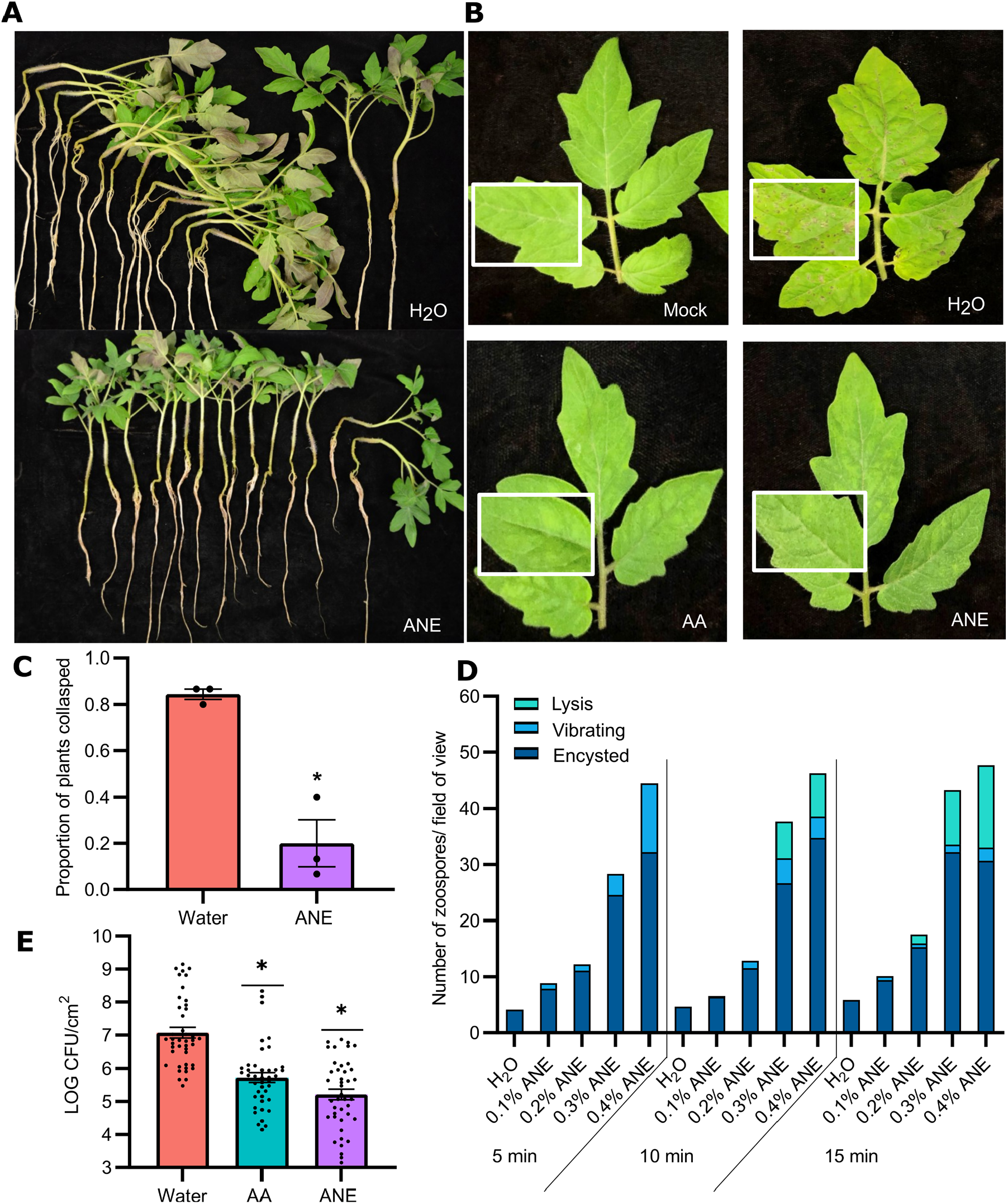
**(A)** Hydroponically grown tomato seedlings treated with 0.4% Acadian (ANE) or water 48 hours post inoculation with *Phytophthora capsici* zoospore suspension and rated on the incidence of collapse at the crown. **(C)** Data are the means and SE for 3 independent trials at 0.4% ANE with 15 plants per treatment per trial. * Significantly different by Wilcoxon rank sums test, *X*^2^ = 3.97, P< 0.046. **(B)** Representative leaf symptoms on hydroponically grown tomato root treated with 0.4% ANE, 10uM Arachidonic Acid (AA), or water 72 hours post spray inoculation with *Pseudomonas syringae* pv. *tomato* bacterial suspension in 10 mM MgCl_2_ at OD^600^= 0.3. Bacterial colonization as measured by LOG colony forming units (CFU) per cm^2^ of leaf tissue72 hours post-inoculation **(E)**. Data are the means and SE for 3 independent trials with n=14 per trial. * Significantly different by Tukey’s HSD *P*<0.0001. **(D)** Direct effect of 0.1%, 0.2%, 0.3% and 0.4% ANE on zoospore motility at 5, 10, and 15 minutes of exposure. The number of lysed, vibrating, and encysted zoospores per field of view. Data are the means for 3 independent trials with n=3 per treatment concentration and timepoint per trial.

**Figure 7.**
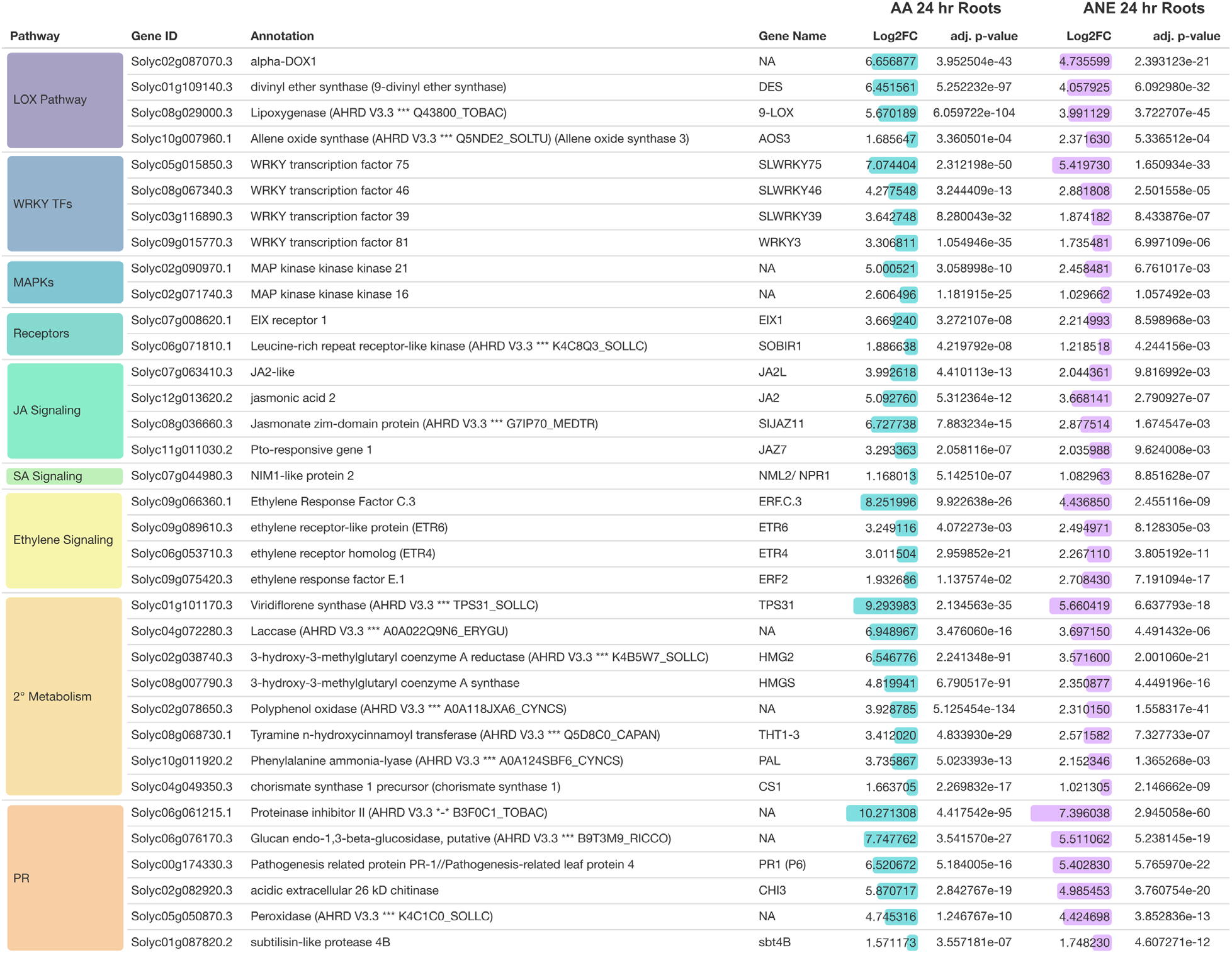
Significantly up-regulated genes shared by Arachidonic Acid (AA) and Acadian (ANE) treated roots at 24 hours. Log_2_-fold change and adjusted *P*-values shown of all genes.

The leaves of plants with roots treated with AA, ANE, or H_2_O were challenged with the bacterial pathogen *Pst*. Both AA- and ANE-root treatments induced systemic resistance to *Pst*. The leaves of seedlings that had been root-treated with either AA or ANE showed significantly reduced bacterial colonization 72 hpi compared to H_2_O-treated/inoculated control seedlings (**Fig. 6B**), with leaf symptoms corresponding to treatment effects on colonization (**Fig. 6E**). H_2_O control plants showed an average bacterial titer of 7.06 log CFU/cm^2^, with AA- and ANE-treated plants showing a 1.25- and 1.35-fold decrease in bacterial growth, respectively. The observed leaf symptoms 72 hpi were also consistent with differences in bacterial colonization. These experiments demonstrate that both AA and ANE induced local and systemic resistance to subsequent infection with an oomycete pathogen after root inoculation or a bacterial pathogen after leaf inoculation.

### Direct effect of ANE on plant growth and zoospore motility

A tradeoff often occurs between plant growth and defense, which is frequently observed in seedlings after treatment with high concentrations of MAMPs, resulting in seedling growth inhibition (Gómez-Gómez et al., 1999). Therefore, we investigated the effect of ANE on plant growth in a hydroponic system. Direct treatment of tomato roots with 0.4% ANE significantly reduced fresh weight biomass compared to water at 72 hours post treatment, consistent with an ANE-associated growth penalty (**Sup. Fig. 4A**). The ANE growth penalty is observed locally in treated roots (**Sup. Fig. 4B**) and distally in shoots (**Sup. Fig. 4C**), consistent with ANE’s ability to systemically alter the transcriptional profile and induce resistance.

Typically, MAMPs are thought to primarily act to inhibit pathogen proliferation through direct perception and defense activation in the plant. However, ANE is a complex mixture with many potentially bioactive compounds. Because a potential direct effect of ANE on zoospores of *Phytophthora* has not been reported, we investigated ANE’s effect on zoospore integrity and encystment. In a concentration-dependent manner, zoospores of *P. capsici* encyst and lyse in the presence of ANE (**Fig. 6D**). Zoospores exposed to dilute ANE (≤ 0.1%) showed abnormal motility or premature encystment compared to water controls, while zoospores treated with ≥ 0.3% ANE showed premature enystment and lysis compared to water controls (**Sup. Table 4**). These data demonstrate ANE’s ability to alter zoospore motility by inducing abnormal movement, encystment, and lysis. Therefore, components in ANE not only trigger defense in tomato, but have the capacity to affect *P. capsici* zoospore behavior and viability following direct exposure.

## Discussion

AA and related eicosapolyenoic fatty acids are unusual elicitors of defense whose structural requirements for activity, absence from higher plants, and abundance in oomycete pathogens distinguish them as MAMPs (Robinson and Bostock, 2015). *A. nodosum*, the brown alga from which ANE is derived, belongs to the same major lineage as oomycetes and contains AA as a predominant polyunsaturated fatty acid (van Ginneken et al., 2011). ANE is used commercially in crops as a biostimulant and may also help plants cope with biotic and abiotic stresses. Through comparative transcriptomic analysis, our study revealed that root treatment with AA or ANE locally and systemically induce similar yet distinct transcriptional profiles in tomato. Root treatment with AA or ANE alter the accumulation of defense-related phytohormones locally in treated roots and systemically in untreated leaves. This study also revealed the systemic nature of AA-induced resistance, and the local and systemic nature of ANE-induced resistance in tomato against pathogens with different parasitic strategies.

Unlike canonical MAMPs that are perceived at the cell surface, AA is rapidly taken up by plant cells and metabolized, with significant incorporation into plant lipids (Preisig and Kuć, 1988; Ricker and Bostock, 1992). Therefore, perception of AA, and by inference the AA present in ANE, is likely different or more complex than direct immune-receptor mediated MAMP recognition and signal transduction. AA can directly engage endogenous plant oxylipin metabolism via action of specific lipoxygenases (LOX) that use AA as a substrate (Andreou et al., 2009; Fournier et al., 1993; Göbel et al., 2002; Göbel et al., 2001; Hwang and Hwang, 2010; Véronési et al., 1996). This study demonstrates AA and ANE induce multiple overlapping local and systemic responses, with interesting parallels and key differences with canonical MAMPs.

AA and ANE locally and systemically alter transcriptional profiles of tomato with many shared and unique features. Varying levels of transcriptional overlap were seen across time points and tissues with up to 80% overlap in roots and up to 55% overlap in leaves between genes differentially expressed compared to water in AA- and ANE-treated plants (**Fig. 3C-D**). Similarly, the canonical MAMPs elf18 and flg22 induce distinct yet primarily overlapping transcriptional changes in *A. thaliana* (Wan et al., 2019). More recent work in *A. thaliana* compared the early transcriptional response of plants treated with diverse MAMPs and damage associated molecular patterns (DAMPs), which elicited striking levels of transcriptional congruency at early time points (5 min to 3 hours) (Bjornson et al., 2021). Like traditional MAMPs, AA and ANE also induce expression of genes associated with PTI including several *WRKY* transcription factors and SA receptor *NPR1*, which also accumulates in response to flg22 treatment in *A. thaliana* (Bjornson et al., 2021; Chen et al., 2017). Despite being an “orphaned” MAMP that may have a different mode of perception, AA and by inference ANE still engage common transcriptional and hormone-mediated defenses.

Systemic resistance is often induced in plants treated with MAMPs and thus is considered a product of the immune response (Mishina and Zeier, 2007). Root treatment with either AA or ANE protected plants locally from *P. capsici*, and systemically from *Pst*. MAMP treatment is also commonly associated with plant growth inhibition due to a growth vs. defense tradeoff (Wang and Wang, 2014). As plants prime defense, there can be down-regulation of photosynthesis-related genes and a shift of photoassimilate from growth to defense, resulting in a growth penalty. Flg22 and elf18 treatment of *A. thaliana* inhibits seedling growth (Gómez-Gómez et al., 1999). Likewise, AA at concentrations used to induce immunity can significantly reduce tomato seedling length and inhibit lateral root formation and cotyledon expansion (Dye and Bostock, 2021). We found that tomato roots treated with 0.4% ANE in a hydroponic system display a significant reduction in root and shoot fresh weight (**Sup. Fig 3**). Our findings coincide with a systemic induced resistance phenotype and growth penalty associated with other well-characterized MAMPs.

Although there is striking overlap between AA/ANE and some MAMP-induced immune responses, there are also distinct differences in how AA/ANE potentially interact with immune signaling and defense. A previous gene expression study in tomato revealed AA-root treatment strongly induces local and systemic expression of several key oxylipin pathway genes (Dye, 2020). Here we show upregulation of the same subset of genes in response to ANE root treatment. Like AA, ANE also activates expression of *α-DOX1* and *9-LOX*, both of which form fatty acid hydroperoxides representing a first step in the generation of plant oxylipins. Oxylipins can serve as signaling molecules to mediate plant responses to wounding, abiotic stress, and pathogen attack (Robinson and Bostock, 2015). As with AA, ANE also induces expression of 9-DES, which can produce novel antimicrobial divinyl ethers that may operate to help restrict *Phytophthora* infections (Weber et al., 1999). Orthologs of *9-LOX* and *9-DES* are present in pepper, potato, and tobacco, and the 9-LOXs in these species can use AA as a substrate (Andreou et al., 2009; Fournier et al., 1993; Göbel et al., 2002; Göbel et al., 2001; Hwang and Hwang, 2010; Véronési et al., 1996). Like AA, ANE induces expression of AOS3, which produces unstable allene oxides from 13-hydroperoxy fatty acids, the first committed step in JA biosynthesis. Previous work with an *aos* mutant in *A. thaliana* established that an intact JA pathway was required for AA-induced resistance to *Botrytis cinerea* (Savchenko et al., 2010). The same study showed that AA treatment of *Arabidopsis* and tomato leaves increased JA and reduced SA levels in the plants, a treatment effect abolished in the case of the *Arabidopsis aos* mutant. These data highlight the critical role of oxylipin metabolism and potentially of oxidized products of AA to help trigger changes in defense hormone signaling.

In the present study, we found accumulation of OPDA, JA, and SA concurrently in AA- and ANE-root treated tomato plants with an induced resistance phenotype. Similarly, Lal et al found that *A. thaliana* with phosphomimetic mutations in receptor-like kinase, BIK1, displayed elevated levels of both SA and JA in noninfected and in *Pst*-challenged plants (Lal et al., 2018). Our data suggest that modulation of JA and SA, classically antagonistic in induced resistance studies, is complex and nuanced in tomato in response to AA and ANE. These findings also suggest that AA and ANE can induce broad-spectrum resistance to pathogens that utilize different parasitic strategies.

While AA and ANE treatment have similar transcriptional outcomes *in planta* and share ability to induce local and systemic resistance, ANE is a complex extract containing EP as well as additional potentially bioactive compounds. Early work with AA demonstrated that it has no direct effect on zoospore motility or viability of *P. infestans* and *P. capsici* (Ricker and Bostock, 1994). In contrast, we found that direct exposure of *P. capsici* zoospores to ANE diminishes motility and viability in a concentration-dependent manner (**Sup. Fig 4, Supp. Table 4**). However, our induced resistance experimental format with *P. capsici* ensured that zoospores did not come into direct contact with inhibitory concentrations of ANE. Also, the relevance of our observation in field settings is unclear since we would expect there to be substantial dilution of ANE during soil applications. Nonetheless, the potential to directly inhibit pathogen inoculum should be considered in experimental design in assessments of seaweed-derived and other biostimulants.

This study provides in-depth profiles of AA- and ANE-associated local and systemic transcriptional remodeling events, phytohormone changes, and induced resistance in tomato with interesting parallels and differences with canonical MAMP action. Further investigation and functional analyses of oxylipin metabolism genes in relation to AA/ANE action is needed to help elucidate their potential role in MAMP signaling. Future research with EP-containing biostimulants will lead to a more holistic understanding of diverse MAMP perception and response, with potential practical implications for crop disease management.

## Materials and Methods

### Disease and plant growth assays

#### Plant Materials and hydroponic growth system

Seeds of tomato (*Solanum lycopersicum cv*. ‘New Yorker’) were surface sterilized and germinated for 10 days on germination paper. Seedlings were transferred to a hydroponic growth system and maintained in a growth chamber as previously described in (Dye, 2020). Seed was obtained from a commercial source (Totally Tomatoes, Randolph, WI).

#### Root treatments

Fatty acid sodium salts (Na-FA; Nu-Chek Prep, Elysian, MN) were prepared and stored as previously described (Dye, 2020). A proprietary formulation of *A. nodosum* extract (ANE; APH-1011-00, Acadian Seaplants, Ltd, Nova Scotia, Canada) was diluted with deionized water (diH_2_O) to a 10% working concentration, which was used to prepare treatment dilutions. All chemicals were diluted to their treatment concentrations with sterile diH_2_O. Hydroponically reared, 3-week-old tomato seedlings with two fully expanded true leaves were transferred to 1 L darkened treatment containers. For *P. capsici* disease assays and ANE growth penalty assessments, roots were treated with an aerated 0.4% ANE solution, or sterile diH_2_O (control) for 72 hours. For *Pst* disease assays, roots were treated with an aerated suspension of 10 µM Na-AA, 0.4% ANE, or diH_2_O for 72 hours. Roots were soaked and rinsed as previously described (Dye and Bostock, 2021). Treated seedlings were then returned to treatment containers with aerated 0.5X Hoagland’s solution for 72 hours followed by harvest for biomass measurements or inoculation with either *P. capsici* or *Pst*.

#### Phytophthora capsici root inoculation and disease assessment

For root/crown disease assays, *P. capsici* isolate PWB-53 (Hensel) was used for inoculation. Tomato seedlings were individually inoculated with 5 mL of zoospore suspension (0.5×10^4^ per mL) as described in (Dye, 2020). Seventy-two hours post inoculation (hpi), disease incidence was rated on the basis of seedling collapse at the crown. Pathogen cultures were maintained, inoculum prepared, and seedling collapse determined as previously described (Dileo et al., 2010; Dye, 2020; Dye and Bostock, 2021).

#### Pst inoculation and analysis of bacterial growth

For leaf disease assays, *Pst* strain DC3000 was used for inoculation. *Pst* from -80°C-maintained glycerol stock was grown on NYGA media amended with rifampicin at 100 µg/mL for two days at 28°C. *Pst* was restreaked on rifampicin-amended NYGA media and grown for 24 hours at 28°C. The bacteria were harvested and resuspended in 5 mM MgCl_2_. Leaves were sprayed with bacterial suspension of OD^600^ = 0.3 with 0.01% Silwett using a Preval spray system, Nokoma Products, Bridgefield, IL). Plants were sprayed until runoff with abaxial and adaxial leaf surfaces covered. Cut Parafilm was used as a protective barrier around the base of plants to prevent contamination of the hydroponic system. Plants were covered with clear plastic bags for 48 hpi. Bacterial colonization was quantified by growth curve analysis 4 days post-inoculation (dpi) as described by (Liu et al., 2009).

### ANE growth penalty assay

Post treatment and Hoagland’s solution interval, tomato seedling were harvested and roots excised from shoots. Root and shoot tissue samples were individually weighed and fresh weights recorded. Roots represent all below surface plant tissue beneath the hypocotyl and shoots represent all aerial tissue including leaves.

### 3’Batch Tag Sequencing Assay

#### Root treatment, tissue harvest, and RNA Extraction

For tissue samples for 3’ Batch Tag Sequencing, roots were treated with an aerated solution of 10µM AA, 10µM LA, 0.4% Acadian, or diH_2_O. Harvested tissue was then subjected to total RNA extraction using Qiagen’s RNeasy Plant Mini Kit, with off column DNase digestion using Qiagen’s RNase-Free DNase Set (Qiagen, Germantown, MD). Each sample was the pool of roots or leaves of two seedlings with three replications per tissue, treatment, and timepoint. All samples were then submitted to the UC Davis Genome Center’s DNA Technology Core for quality control via bioanalyzer analysis, RNA-seq library generation, and sequencing. Gene expression profiling was carried out using a 3’-Tag-RNA-Seq protocol. Barcoded sequencing libraries were prepared using the QuantSeq FWD kit (Lexogen, Vienna, Austria) for multiplexed sequencing according to the recommendations of the manufacturer using both the UDI-adapter and UMI Second-Strand Synthesis modules (Lexogen). The fragment size distribution of the libraries was verified via micro-capillary gel electrophoresis on a LabChip GX system (PerkinElmer, Waltham, MA). The libraries were quantified by fluorometry on a Qubit fluorometer (LifeTechnologies, Carlsbad, CA), and pooled in equimolar ratios. The library pool was Exonuclease VII (NEB, Ipswich, MA) treated, SPRI-bead purified with KapaPure beads (Kapa Biosystems / Roche, Basel, Switzerland), and quantified via qPCR with a Kapa Library Quant kit (Kapa Biosystems) on a QuantStudio 5 RT-PCR system (Applied Biosystems, Foster City, CA). Up to forty-eight libraries were sequenced per lane on a HiSeq 4000 sequencer (Illumina, San Diego, CA) with single-end 100 bp reads.

#### RNAseq data processing and analysis

The raw reads were imported into the Galaxy platform for comprehensive data analysis including quality control, alignment, and differential expression analysis (Goecks et al., 2010). The raw data was processed using the quality control tool FastQC/MultiQC (v1.7) to access the quality of the raw sequence data (Ewels et al., 2016). Read alignment was conducted using RNA STAR aligner (v2.6.0b-1) and post alignment quality control employed FastQC/MultiQC (v1.7) (Ewels et al., 2016). Quantification of reads per gene was carried out using featureCounts (v1.6.3) (Liao et al., 2013). Read counts were normalized and differential gene expression analysis conducted using DESeq2 (v2.11.40.6) (Love et al., 2014). Differential genes were visualized using the Heatmap2 (v1.0) program followed by functional enrichment of differential genes by the program GoSeq (1.34.0) (Young et al., 2010). All aforementioned bioinformatics programs were accessed through the Galaxy toolshed (Blankenberg et al., 2014). Differential gene expression was also visualized as volcano plots developed using ggplot2 (v3.3.5) via a custom script (Supplemental_File_1).

### Phytohormone quantitation

For phytohormone quantitation, roots were treated with an aerated suspension of 10 µM Na-AA (in deionized water), 0.4% ANE (in deionized water), or deionized water. Following 24- and 48-hours root exposure to their respective treatments, plants were harvested, root and leaf tissue dissected from shoots, and the collected tissue flash frozen in liquid nitrogen. Root and leaf tissue samples were submitted to the Donald Danforth Plant Science Center’s Proteomics and Mass Spectrometry Facility for acidic plant hormone extraction and quantification. Each sample was the pool of roots and leaves of three seedlings with three samples per tissue, treatment, and timepoint. The experiment was performed once.

Analytical reference standards were used for the following analytes: indole-3-acetic acid (IAA; Sigma-Aldrich St. Louis, MO), N-(3-indolylacetyl)-DL-aspartic acid (IAA-Asp; Sigma-Aldrich, St. Louis, MO), (+/-)-jasmonic acid (JA; Tokyo Chemical Industry Company, Tokyo, Japan), salicylic acid (SA; Acros Organics, Geel, Belgium), (+/-)-abscisic acid (ABA; Sigma-Aldrich, St. Louis, MO), N-jasmonyl-L-isoleucine (JA-Ile; Toronto Research Chemicals, Toronto, ON), 12-oxo-phytodienoic acid (OPDA; Cayman Chemical, Kalamazoo, MI), *cis*-zeatin (*c*Z; Santa Cruz Biotechnology, Santa Cruz, CA), *trans*-zeatin (*t*Z; Caisson Labs, Smithfield, UT), DL-dihydrozeatin (DHZ; Research Products International, Mount Prospect, IL), and *trans*-zeatin riboside (*t*ZR; Gold Biotechnology, St. Louis, MO). Internal standards were d_5_-JA (Tokyo Chemical Industry Company, Tokyo, Japan), d_5_-IAA (CDN Isotopes, Pointe-Claire, QC), d_5_-dinor-OPDA (Cayman Chemical, Kalamazoo, MI), d_6_-SA(CDN Isotopes, Pointe-Claire, QC), d_6_-ABA (ICON Isotopes, Dexter, MI), d_5_-*trans*-zeatin (OlChemIm, Olomouc, Czech Republic), d_5_-*trans*-zeatin riboside (OlChemIm, Olomouc, Czech Republic), ^13^C_6_^15^N JA-Ile (New England Peptide, Gardner, MA), and ^13^C_4_^15^N IAA-Asp (New England Peptide, Gardner, MA). LC-MS grade methanol (MeOH) and acetonitrile (ACN) were sourced from J.T. Baker (Avantor Performance Materials, Radnor, PA) and LC-MS grade water was purchased from Honeywell Research Chemicals (Mexico City, Mexico). Standard and internal standard stock solutions were prepared in 50% methanol and stored at -80° C. Calibration standard solutions were prepared fresh in 30% methanol.

#### Phytohormone Extraction

Phytohormones *c*/*t*Z, DHZ, *t*ZR, SA, ABA, IAA, IAA-Asp, JA, JA-Ile and OPDA were extracted at a tissue concentration of 100 mg/mL in ice cold 1:1 MeOH: ACN. Around 100 mg of tissue sample were weighed and 10 µL of mixed stable-isotope labeled standards (1.0 μM for d_5_-*t*Z and d_5_-*t*ZR, 2.5 μM for d_4_-SA, d_6_-ABA, d_5_-JA, d5-IAA, ^13^C_6_^15^N-IAA-Asp and d5-dinor-OPDA, and 25.0 μM for ^13^C_6_^15^N-JA-Ile) were added to each sample prior to extraction. The samples were homogenized with a TissueLyzer-II (Qiagen) for 5 minutes at 15 Hz and then centrifuged at 16,000 x g for 5 minutes at 4° C. The supernatants were transferred to new 2 mL tubes and the pellets were re-extracted with 600 µL 1:1 ice cold MeOH: ACN. These extracts were combined and dried in a vacuum centrifuge. The samples were then reconstituted in 100 µL 30% methanol, centrifuged to remove particulates, and then passed through a 0.8 μm polyethersulfone spin filter (Sartorius, Stonehouse, UK) prior to dispensing into HPLC vials for LC-MS/MS analysis.

#### LC-MS/MS Analysis

Phytohormones *c*/*t*Z, DHZ, *t*ZR, SA, ABA, IAA, IAA-Asp, JA, JA-Ile and OPDA were quantified using a targeted multiple reaction monitoring (MRM)/isotope dilution-based LC-MS/MS method. A Shimadzu Prominence-XR UFLC (UPLC) system connected to a SCIEX hybrid triple quadrupole-linear ion trap MS equipped with Turbo V^™^ electrospray ionization (ESI) source (SCIEX, Framingham, MA) were used for the quantitative analysis. Ten microliters of the reconstituted samples were loaded onto a 3.0 × 100 mm 1.8 μm ZORBAX Eclipse XDB-C_18_ column (Agilent Technologies, Santa Clara, CA, USA) and the phytohormones were eluted within 22.0 minutes, in a binary gradient of 0.1% acetic acid in water (mobile phase A) and 0.1% acetic acid in 3:1 ACN:MeOH (mobile phase B). The initial condition of the gradient was 5% B from 0 to 2.0 minutes, ramped to 40% B at 10.0 minutes, further ramped to 50%B at 15.0 minutes, and then quickly raised to 95% B at 19.0 minutes and kept at 95% B until 22.0 minutes. The flow rate was set at 0.4 mL/min. Source parameters were set as follows: curtain gas 25 psi; source gas 140 psi; source gas 250 psi; collisionally activated dissociation (CAD) gas set to ‘medium’; interface heater temperature 500°C; ion spray voltage set to +5500 V for positive ion mode and -4500 V for negative ion mode. Individual analyte and internal standard ions were monitored using previously optimized MRM settings programmed into a polarity switching method (cytokinins and auxins detected in positive ion mode, others detected in negative ion mode). Analyst 1.6.2 software (SCIEX, Framingham, MA) was used for data acquisition; MultiQuant 3.0.2 software (SCIEX, Framingham, MA) was used for data analysis. The detected phytohormones were quantified based upon comparison of the analyte-to-internal standard integrated area ratios with a standard curve constructed using those same analytes, internal standards and internal standard concentrations (2.5 μM ^13^C_6_^15^N-JA-Ile; 0.10 μM d_5_-*t*Z and d_5_-*t*ZR; others 0.25 μM). The mixed calibration solutions were prepared over the range from 1.0 fmol to 100 pmol loaded on the column. The actual calibration range for each analyte was determined according to the concentrations of the analyte in samples.

### ANE zoospore motility assay

Aliquots of *P. capsici* zoospore suspension at 10^6^ zoospores/mL were distributed to a polystyrene ninety-six well plate (Fisher Scientific, Hampton, NH) and exposed to ANE such that the final concentrations per well were 0.1%, 0.2%, 0.3%, and 0.4% ANE. Sterile deionized water was used as a negative control. At 5, 10, and 15 minutes of exposure to their respective treatments, a hemocytometer was used to quantify vibrating and encysted zoospores per field of view. An overall motility status was also observed where fields of view with no motile zoospores remaining were reported. Using the standardized starting concentration, the overall motility status of the replicate, and the sum of encysted and vibrating zoospores and the number of lysed zoospores was calculated. Pathogen cultures were maintained and zoospore suspensions prepared as previously described (Dye, 2020)

## Acknowledgements

Research supported in part by a Gates Millenium Scholars Program fellowship and Jastro-Shields awards to DCL, United States Department of Agriculture National Institute of Food and Agriculture grant 2021-67034-35049 to DMS, an NIH grant R35GM136402 to GC, and an unrestricted gift from Acadian Seaplants LTD to RMB. Sequencing was performed by the DNA Technologies and Expression Analysis Core at the UC Davis Genome Center, supported by NIH Shared Instrumentation Grant 1S10OD010786-01. Phytohormone analyses were performed by the Donald Danforth Plant Science Center, St. Louis, MO USA, based upon work supported by the National Science Foundation under Grant No. DBI-1427621 for acquisition of the QTRAP LC-MS/MS.

## Author Contributions

Study planned by all authors. Experiments performed by DCL and analyzed by DCL, DMS, and RMB. Paper written by DCL and RMB, with editing by DMS and GLC.

## Notes

### Competing Interest Statement

The authors have declared no competing interest.

