## Supplemental Figures for "Overlapping local and systemic defense induced by an oomycete fatty acid MAMP and brown seaweed extract in tomato"

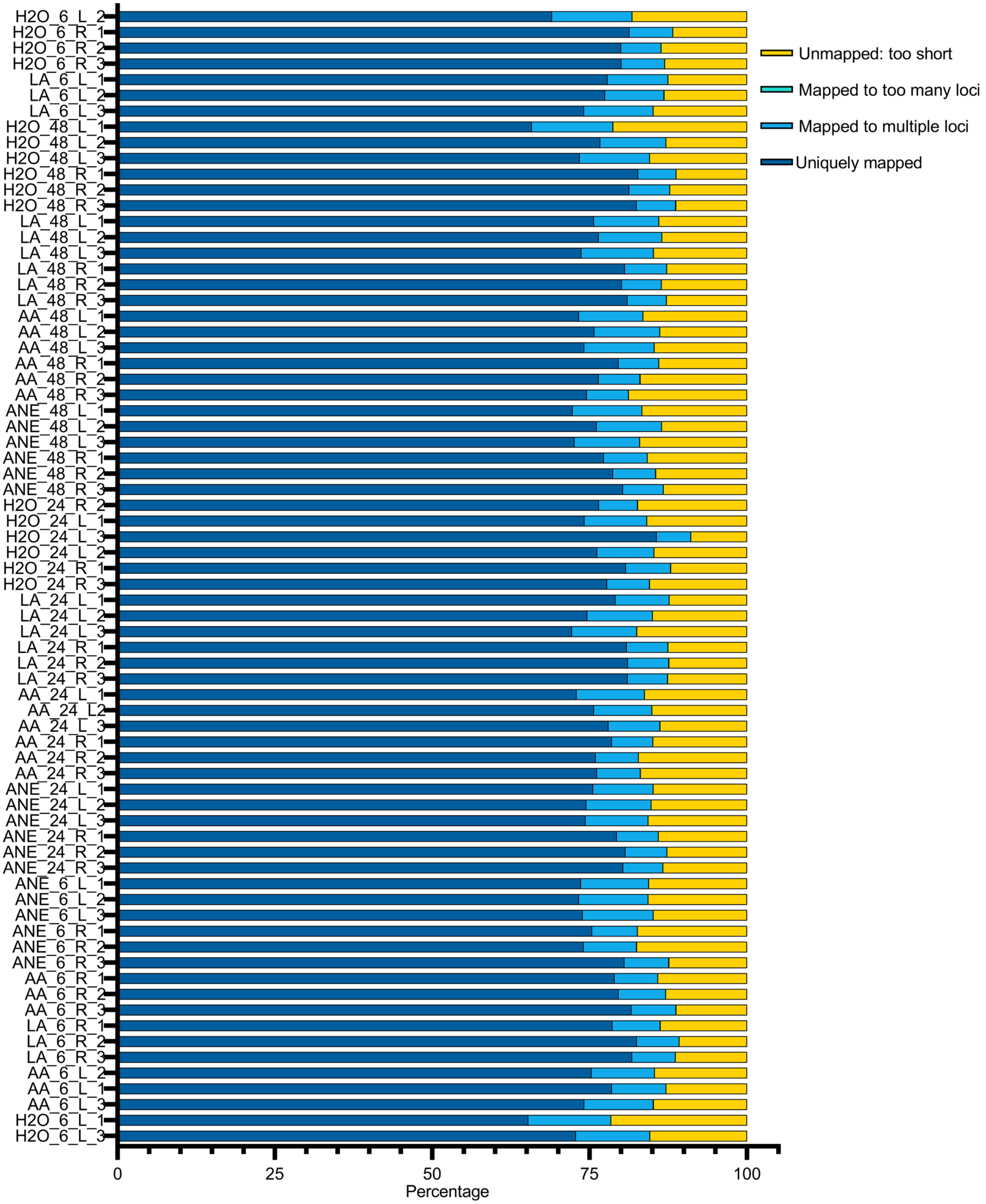


**Supplementary Figure 1.** Percent of sequencing reads that uniquely mapped, mapped to multiple loci, mapped to many loci, and too short unmapped reads to the tomato genome build SL 3.0. Tomato roots were treated with 10 µM Arachidonic Acid (AA), 0.4% Acadian (ANE), $H_2O$, or 10 µM Linoleic acid (LA). Following 6, 24, and 48 hours root exposure to their respective treatments, plants were harvested, root (R) and leaf (L) samples were processed and subjected to RNA sequencing. Alignment was conducted using RNA STAR aligner accessed through the Galaxy toolshed.

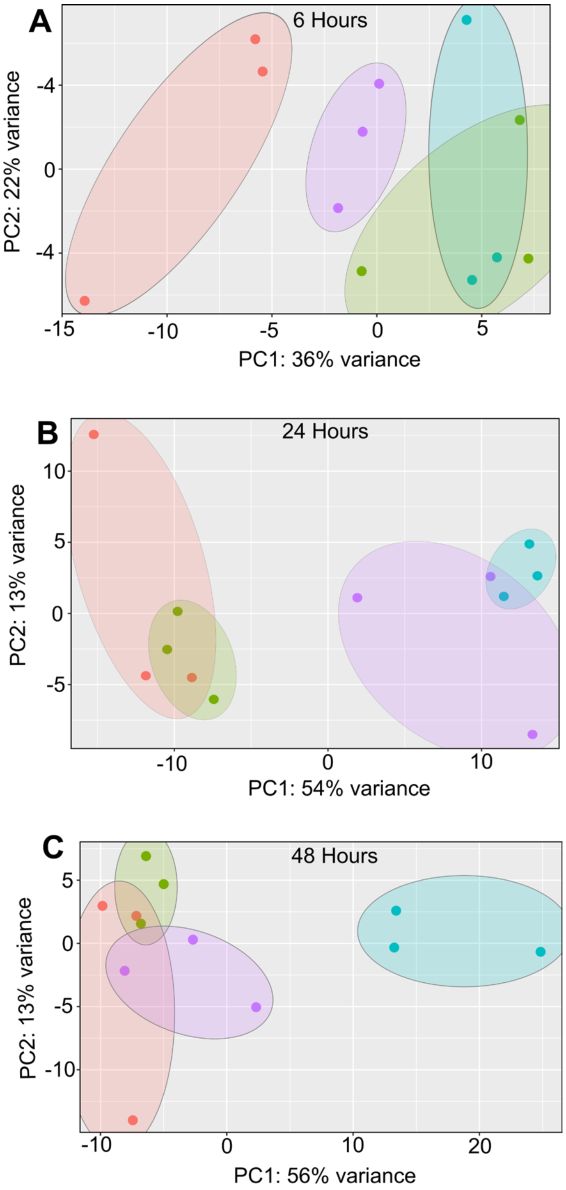


**Supplemental Figure 2.** Principle component analysis scatterplots of RNA sequencing data in leaves after **(A)** 6, **(B)** 24, and **(C)** 48 hours of treatment with 10 $\mu$M Arachidonic Acid (AA), 0.4% Acadian (ANE), $H_2O$, or 10 $\mu$M Linoleic acid (LA). PCA was conducted using the normalized read counts for all samples. PCA plots show variance of three biological replicates performed per timepoint and treatment.

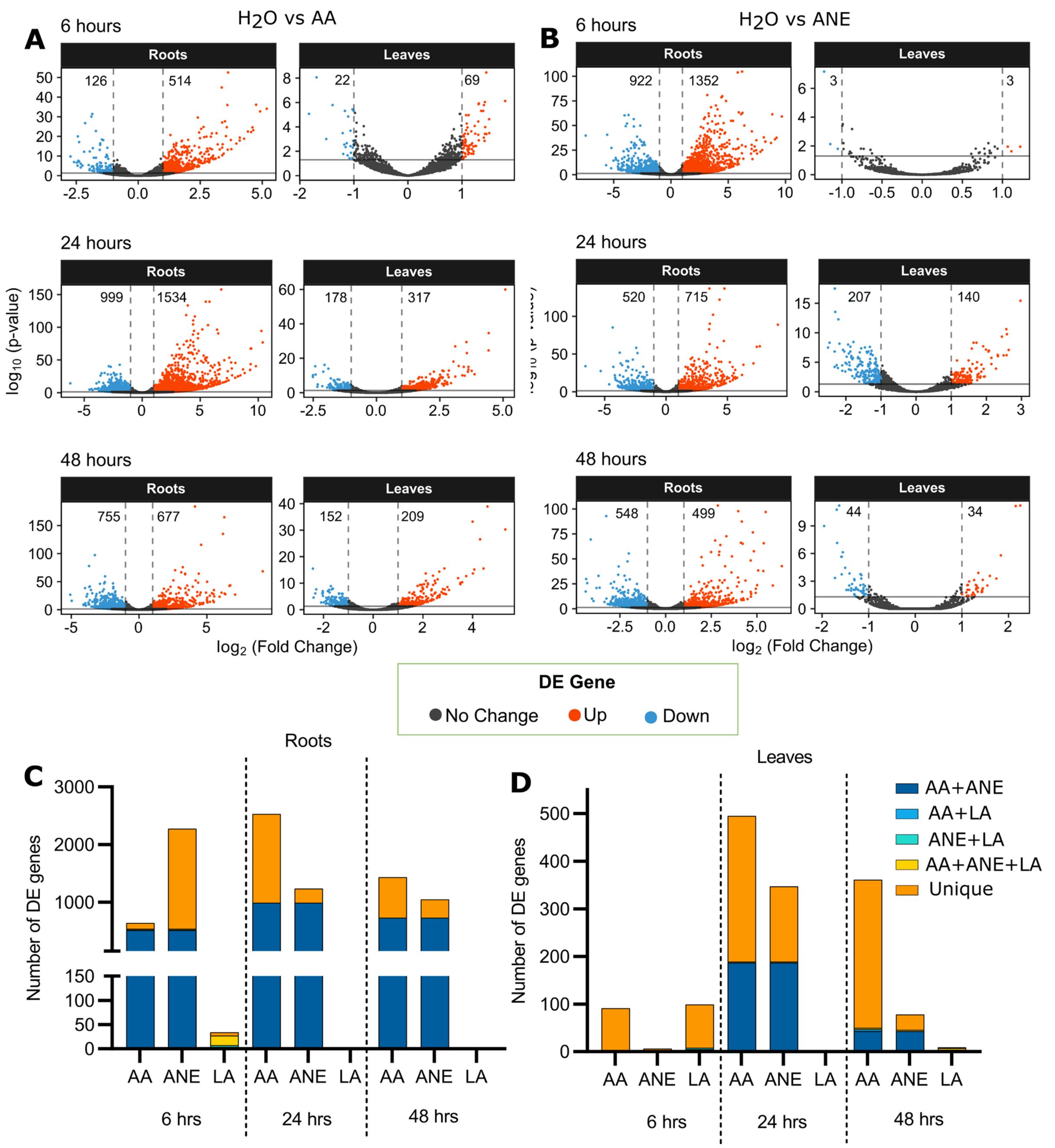


**Supplemental Figure 3.** Scatter plot of DE genes with the number of significantly up (red) and down (blue) regulated genes plotted from roots and leaves treated with AA **(A)** or ANE **(B)** compared to $H_2O$-treated tissue across time points. Solid and dashed lines represent cutoffs for significant DE genes (adjusted *P*-value < 0.05 and an absolute fold change in gene expression >2). Number of DE genes shared and unique between AA, ANE, and Linoleic acid (LA) compared to $H_2O$ in **(C)** roots and **(D)** leaves. Plant roots were treated with $H_2O$, 10 µM LA, 10 µM AA, or 0.4% ANE for 6, 24 or 48 hours. DE genes compared to $H_2O$ have an adjusted *P*-value < 0.05 and an absolute fold change in gene expression >2.

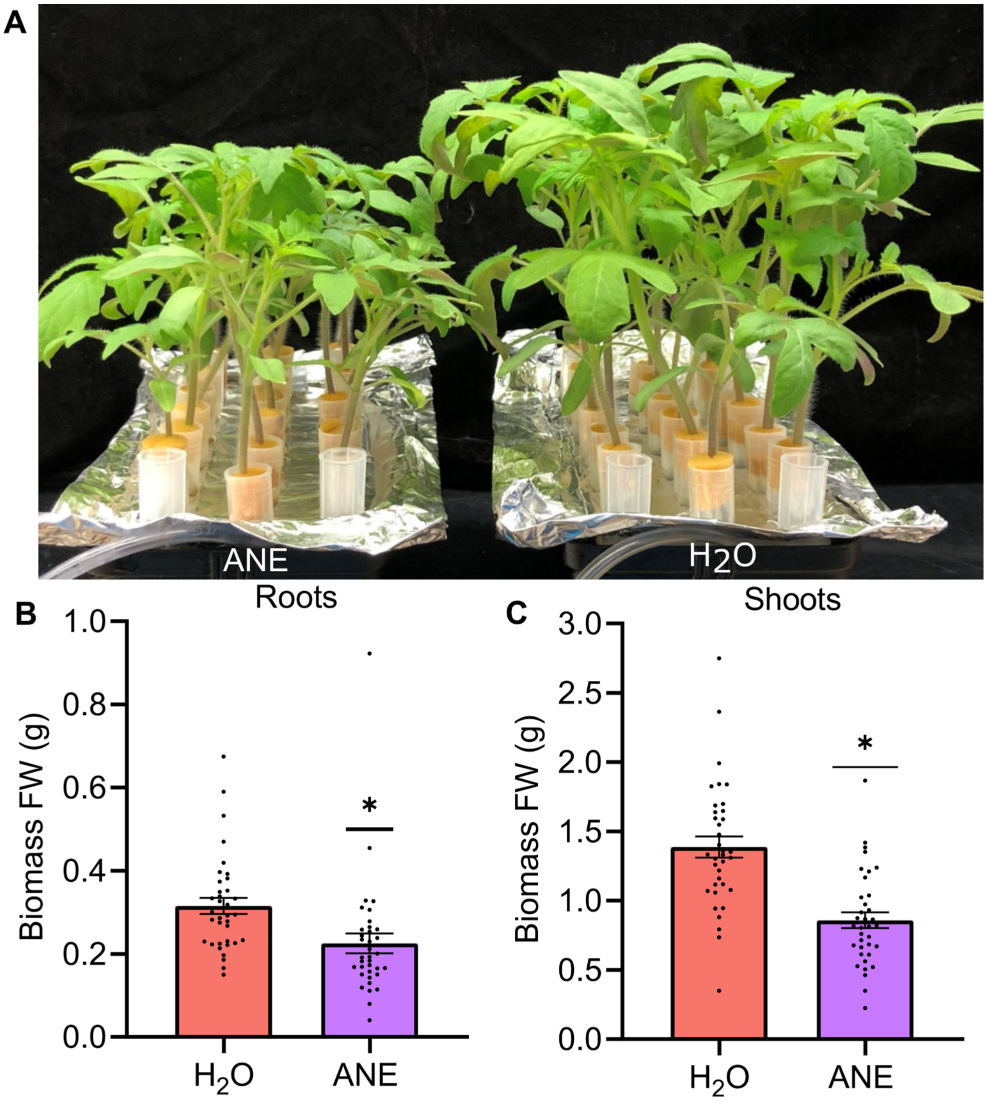


**Supplemental Figure 4. (A)** Acadian (ANE)-associated growth penalty in hydroponically-reared tomato 72 hours post treatment with 0.4% ANE. Treatment effect on fresh weight biomass of **(B)** roots and **(C)** shoots. Data are the means and SE for 2 independent trials at 0.4% ANE with 18 plants per treatment per trial. * Significantly different by Student's t test, $P<$ 0.0048 (Panel B) and $P<0.0001$ (Panel C). Error bars represent standard error of 2 biological replicates.
